# Stray cats and dogs carrying zoonotic *Enterocytozoon bieneusi* genotype D in China: a public health concern

**DOI:** 10.1101/2023.04.17.537122

**Authors:** Yidan Zhang, Yan Zhang, Rongsheng Mi, Luming Xia, Hongxiao Han, Tao Ma, Haiyan Gong, Yan Huang, Xiangan Han, Zhaoguo Chen

**Author notes:** Yan Zhang and Yidan Zhang contributed equally to this work. Corresponding author* (ZGC).

## Abstract

*Enterocytozoon bieneusi* is reported to be a common microsporidian of humans and animals in various countries. However, scarce information on *E. bieneusi* has been recorded in cats (*Felis catus*) and dogs (*Canis familiaris*) in China. As such, we undertook molecular epidemiological investigation of *E. bieneusi* in cats and dogs in Shanghai, China. A total of 359 genomic DNAs were extracted from individual faecal samples from cats (*n* = 59) and dogs (*n* = 300), and then tested using a nested PCR-based sequencing approach employing internal transcribed spacer (ITS) of nuclear ribosomal DNA as the genetic marker. *Enterocytozoon bieneusi* was detected in 34 of all 359 (9.5%) faecal samples from cats (32.2%; 19/59) and dogs (5.0%; 15/300), including 24 stray cats and dogs (22.6%; 24/106), as well as ten household/raised cats and dogs (4.0%; 10/253). Correlation analyses revealed that *E. bieneusi* positive rates were significantly associated with stray cats and dogs (*P* < 0.05). The analysis of ITS sequence data revealed the presentation of five known genotypes CD7, CHN-HD2, D, PtEb IX and Type IV and two novel genotypes D-like1 and PtEb IX-like1. Zoonotic genotype D was the predominant type with percentage of 61.8 (21/34). Phylogenetic analysis of ITS sequence data sets showed that genotypes D, D-like1 and Type IV clustered within Group 1, showing zoonotic potential. The others were assigned into Group 10 with host specificity. These findings suggested that cats and dogs in Shanghai harbor zoonotic genotype D of *E. bieneusi* and may have a significant risk for zoonotic transmission. Further insight into the epidemiology of *E. bieneusi* in animals, water and the environment from other areas in China will be important to have an informed position on the public health significance of microsporidiosis caused by this microbe.

## Introduction

*Enterocytozoon bieneusi* is the commonest pathogen responsible to most of human microsporidiosis, causing chronic or severe diarrhea, malabsorption or wasting [1, 2]. This microbe can transmit through faecal-oral route, via spores contaminated water, food or direct contact with infected individuals or their droppings [3]. Typically, molecular method PCR-based sequencing of internal transcribed spacer (ITS) of ribosomal DNA has been widely used to identify *E. bieneusi* [2]. Using this approach, more than 600 genotypes have been identified in a broad host range [4] (review). Some of these genotypes can be only found in animals (e.g., genotypes SCC-2 [Common chipmunk]), however, many other genotypes have been recorded in both humans and animals showing zoonotic potential (e.g., genotypes EbpC, D and Type IV). Thus, the National Institute of Allergy and Infectious Diseases (NIAID) classifies *E. bieneusi* as a Category B Priority Pathogen [5].

Numerous studies have been investigated *E. bieneusi* from humans and a large group of animal species, including various orders of mammals (Artiodactyla, Carnivora, Diprotodontia, Lagomorpha, Perissodactyla, Primates and Rodentia), birds (Anseriformes, Columbiformes, Falconiformes, Galliformes, Passeriformes, Psittaciformes and Struthioniformes) and reptiles (Squamata) as well as insects (Diptera) in more than 40 countries [4]. Although, there have been > 30 studies investigating this microbe in cats and dogs worldwide [6-8], only ten investigations of *E. bieneusi* was conducted in China, leading the systematic epidemiological studies and risk factors (e.g., temperature and humidity) of *E. bieneusi* in cats and dogs are scarce.

Shanghai is a developed metropolitan city with nearly 25,000 thousand people, and lots of residents in Shanghai have pets (e.g., cats and dogs). Xu et al. studied *E. bieneusi* in cats and dogs in Shanghai with the prevalence of 5.9% [9]. Also, Liu et al. investigated this pathogen from stray dogs (8.8%) and found stray dogs have higher risk to infect humans than pets [10]. Previously, we carried out epidemiological studies of *E. bieneusi* from alpacas [11], cats and dogs [12], farmed cattle [13], farmed goats and sheep [14], wild deer [15], wild marsupials [16], zoo animals [17] and humans [18], in Australia and China. The prevalence and risk factors such as host species, age, sex, location, temperature and season were analysed and *E. bieneusi* genotypes were identified. The results show potential zoonotic transmission and a strong significant association between some risk factors and *E. bieneusi* prevalence. Here, in this study, we investigated *E. bieneusi* in cats and dogs in Shanghai. The aims of this study are to investigate the prevalence of *E. bieneusi* and its risk factors, characterise genotypes and analyze their zoonotic potential.

## Materials and Methods

### Samples and DNA isolation

In total, 359 faecal samples were collected from cats (*Felis catus*) (*n* = 59) and dogs (*Canis familiaris*) (*n* = 300), including household/raised cats (*n* = 9) and dogs (*n* = 244) from pet clinics (*n* = 193) and breeding centers (*n* = 60), as well as stray cats (*n* = 50) and dogs (*n* = 56) in Minhang (*n* = 309) and Jingan (*n* = 50) districts in Shanghai from October 2019 to July 2020, corresponding to three seasons: autumn (*n* = 121), spring (*n* = 128) and summer (*n* = 110) (Table 1). All cats and dogs from pet clinics were maintained in individual cages, while others from breeding centers were raised together. Most of them were apparently healthy. Faecal samples were collected from cats and dogs rectum and most of them were firm and solid, except for a few soft and watery cases. Genomic DNA was extracted directly from 0.1 g to 0.4 g of each of the 359 faecal samples (i.e., right after the sample collection) using the FastDNA SPIN Kit for Soil (MP Biomedicals, Santa Ana, CA, USA).

**Table 1.** The information regarding faecal samples collected from household cats and dogs from pet clinics (*n* = 193) and breeding centers (*n* = 60), as well as stray cats and dogs (*n* = 106) located in Minhang and Jingan districts in Shanghai, China (2019 - 2020).

| Host | Tested | Sample source | Location | *E. bieneusi* prevalence % |
| --- | --- | --- | --- | --- |
| Season | sample nos. | Pet clinic/Breeding center/Stray | Minhang/ Jingan | (test-positive sample nos./total tested sample nos.) |
| **Cat** | **59** | **9/0/50** | **9/50** | **32.2 (19/59)** |
| Spring | - | - | - | - |
| Summer | 50 | 0/0/50 | 0/50 | 38.0 (19/50) |
| Autumn | 9 | 9/0/0 | 9/0 | - |
| **Dog** | **300** | **184/60/56** | **300/0** | **5.0 (15/300)** |
| Spring | 128 | 128/0/0 | 128/0 | 5.5 (7/128) |
| Summer | 60 | 0/60/0 | 60/0 | 5.0 (3/60) |
| Autumn | 112 | 56/0/56 | 112/0 | 4.5 (5/112) |
| **Total** | **359** | **193/60/106** | **309/50** | **9.5 (34/359)** |

### Nested PCR-based sequencing of *E. bieneusi* ITS

Individual genomic DNA samples were subjected to nested PCR-coupled sequencing of the ITS region using an established technique [15]. Briefly, in the first PCR round, primers MSP-1 (forward: 5′-TGA ATG KGT CCC TGT-3′) and MSP-2B (reverse: 5′-GTT CAT TCG CAC TAC T-3′) were used to amplify 601 bp of ITS plus flanking gene sequences. In the second round, primers MSP-3 (forward: 5′-GGA ATT CAC ACC GCC CGT CRY TAT-3′) and MSP- 4B (reverse: 5′-CCA AGC TTA TGC TTA AGT CCA GGG AG-3′) were employed to amplify a product of 535 bp containing 130 bp of the 3′-end of the small subunit (*SSU*) of the nuclear rRNA gene, 243 bp of the ITS and 162 bp of the 5′-region of the large subunit (*LSU*) rRNA gene.

Nested PCR for amplification of ITS was conducted in a reaction volume of 50 μl in a standard buffer containing 4.0 μM MgCl_2,_ 0.4 mM dNTPs, 50 pmol of each primer, 1.25 U of Ex Taq DNA Polymerase (TaKaRa Bio Inc., Beijing, China) and DNA template - except for the negative (no-template) controls. Known test-positive, test-negative and no template controls were included in each PCR run. The cycling conditions for both primary and secondary (nested) PCRs were: 94 °C for 5 min (initial denaturation), followed by 35 cycles of 94 °C for 45 s (denaturation), 54 °C for 45 s (annealing) and 72 °C for 1 min (extension), followed by 72 °C for 10 min (final extension).

The secondary PCR products were examined by gel electrophoresis on a 1.5% agarose gel containing 4S Green Plus Nucleic Acid Stain (Sangon Biotech, Shanghai, China) using TBE (65 mM Tris-HCl, 27 mM boric acid, 1 mM EDTA, pH 9; Bio-Rad, Hercules, CA, USA) as the buffer, and their size estimated using a 2000 bp-DNA ladder (TaKaRa Bio Inc., Beijing, China) as a reference and directly sequenced using primers MSP-3 and MSP-4B in separate reactions. ITS sequences obtained (GenBank accession nos. OQ597705-OQ597711) were inspected for quality using the program Geneious v.10 [19], and compared with reference sequences acquired from the GenBank database (S1 Table). Genotypes of *E. bieneusi* were named according to the recommendations by Santín and Fayer [3, 20].

### Phylogenetic analysis

ITS sequences from this and previous studies were aligned over a consensus length of 301 positions using the methods from Zhang et al. [18], and then subjected to phylogenetic analyses using the Bayesian inference (BI) and Monte Carlo Markov Chain (MCMC) methods in MrBayes v.3.2.3 [21]. The Akaike Information Criteria (AIC) test in jModeltest v.2.1.7 [22] was used to evaluate the likelihood parameters set for BI analysis. Posterior probability (pp) values were calculated by running 2,000,000 generations with four simultaneous tree-building chains, with trees saved every one hundredth generation. A 50% majority rule consensus tree for each analysis was constructed based on the final 75% of trees generated by BI. *Enterocytozoon bieneusi* clades and subclades were assigned using an established classification system [23-27].

### Statistical analysis

The multivariate logistic linear regression were utilised to compare *E. bieneusi* test-positives (faecal samples) with risk factors, and to test the association between the prevalence of *E. bieneusi* DNA and season. The strength of association between *E. bieneusi* prevalence and a univariate risk factor was measured using the odds ratio (OR) calculated with 95% confidence intervals (95% CI). A *P*-value of < 0.05 was considered statistically significant. IBM SPSS Statistics 25.0 (SPSS Inc., Chicago, IL, USA) was used for all statistical analyses [18].

## Results

### Prevalence of *E. bieneusi* and risk factors

*Enterocytozoon* DNA was detected in 34 of the 359 (9.5%) faecal samples from cats (32.2%; 19/59) and dogs (5.0%; 15/300), including 24 stray cats and dogs (19 stray cats and five stray dogs) (22.6%; 24/106), as well as ten household/raised cats and dogs (4.0%; 10/253) (Table 1). None of *E. bieneusi* test-positivity was found in household cats. The prevalences in each season are 20.0% (22/110) in summer, 5.5% (7/128) in spring and 4.1% (5/121) in autumn (Table 2). The association analyses showed that *E. bieneusi* contamination in cats and dogs in summer was higher than that in autumn (OR = 2.841; 95% CI [0.967-8.343]) and spring (OR = 2.117; 95% CI [0.803-5.580]) without significance (*P* > 0.05). There were significant associations of *E. bieneusi*-positivity with living status of cats and dogs (Table 2), stray cats and dogs had 7.112 times higher risk of *E. bieneusi* infection than household/raised cats and dogs (OR = 7.112; 95% CI [3.263-15.500]) (*P* < 0.05).

**Table 2.** Association analysis of the risk factors (seasons and living status) with *Enterocytozoon bieneusi* test-positivity assessed using the multivariate logistic linear regression.

| **Factors** | **Test positive samples nos.** | **Total tested sample nos.** | **Prevalence (%)** | **Odds ratio (95% CI)** | **P-value** |
| --- | --- | --- | --- | --- | --- |
| **Seasons** |  |  |  |  |  |
| Spring | 7 | 128 | 5.5 | 2.117 (0.803-5.580) | 0.152 |
| Summer<sup>1</sup> | 22 | 110 | 20.0 | - | - |
| Autumn | 5 | 121 | 4.1 | 2.841 (0.967-8.343) | 0.075 |
| **Living status** |  |  |  |  |  |
| Stray | 24 | 106 | 22.6 | 7.112 (3.263-15.500) | 0.000* |
| Household/raised | 10 | 253 | 4.0 | - | - |

The strength of association was measured using an odds ratio calculated with 95% confidence intervals (95% CI), and statistical significance was given as a *P*-value. \* = Statistically significant ( $P < 0.05$ ). - = Not available. <sup>1</sup> = Value were used as references when odds ratio was calculated.

### Genotypes and phylogeny

The sequencing of the 301 ITS amplicons (241 - 243 bp) and their subsequent comparisons with reference sequences from the GenBank database revealed that five known genotypes CD7 (1), CHN-HD2 (1), D (21), PtEb IX (7) and Type IV (1) representing 31 amplicons and two novel genotypes D-like1 (2) and PtEb IX-like1 (1) (S2 Table). Genotype D was the most frequent type with the percentage of 61.8% (21/34), followed by genotypes PtEb IX 20.6% (7/34) and D-like1 5.9% (2/34). The rest of genotypes had the same percentage of 2.9% (1/34).

The ITS sequences for all seven genotypes defined herein were aligned with sequences representing all eleven established Groups of *E. bieneusi* [23-27] and then subjected to phylogenetic analysis (Fig 1). In this analysis, All groups were each strongly supported (pp = 0.92 to 1.00) except for Groups 2, 6 and 7 (i.e., pp < 0.85 are not shown). Based on this analysis, genotypes D, D-like1 and Type IV were assigned to Group 1 with strong statistical support (pp = 0.92), others were fall into Group 10 (pp = 0.99).

**Figure 1.**
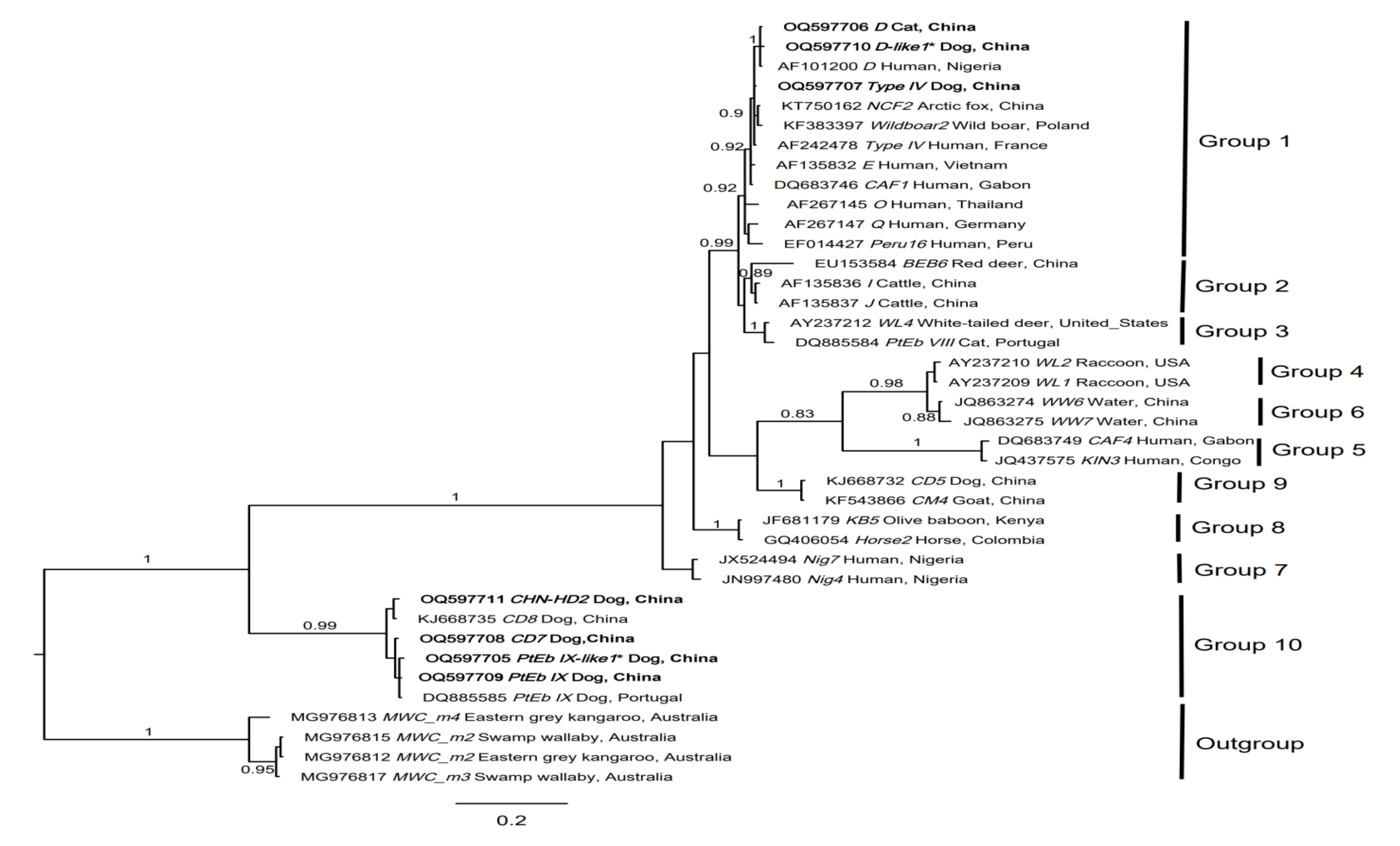
Relationships among genotypes of *Enterocytozoon bieneusi* recorded in cats and dogs in this study inferred from phylogenetic analysis of sequence data for the internal transcribed spacer (ITS) of nuclear ribosomal DNA by Bayesian inference (BI). Sequences from a range of distinct *E. bieneusi* genotypes from published papers were included for comparison in the analysis (S1 Table) [23-27]. Statistically significant posterior probabilities (pp) are indicated on branches. Individual GenBank accession numbers precede genotype designation (in italics), followed by sample and locality descriptions. *Enterocytozoon bieneusi* genotypes identified and characterised from faecal DNA samples in the present study are indicated in bold-type. Clades were assigned group names based on the classification system [23-27]. The scale-bar represents the number of substitutions per site. The *E. bieneusi* genotypes MWC_m2-m4 from marsupials were used as outgroups.

## Discussion

Here, we investigated the distribution and genetic identity of *E. bieneusi* in cats and dogs faecal DNA samples by PCR-based sequencing of ITS in Shanghai, China. In total, 34 of 359 (9.5%) faecal DNA samples were test-positive for *E. bieneusi*, including 19 in cats (32.2%; 19/59) and 15 in dogs (5.0%; 15/300). This is the first time that the highest prevalence of *E. bieneusi* was found in cats around the world, and genotype D - the most frequently identified in humans, was widely recorded in both cats and dogs.

This study revealed a prevalence of *E. bieneusi* of 32.2% (19/59) in cats and 5.0% (15/300) in dogs from pet clinics, breeding centers and shelters in Shanghai, China. The total prevalences of *E. bieneusi* in cats and dogs worldwide are reported to range from 1.4% (2/143) [28] to 31.3% (25/80) [29] and 0.8% (2/237) [6] to 22.9% (149/651) [30], respectively (Table 3). The prevalence of *E. bieneusi* in cats herein is the highest record globally and the *E. bieneusi* test-positives in cats were all stray cats. Whereas, *E. bieneusi* prevalence in dogs is only higher than that recorded in a few *E. bieneusi* studies in dogs (e.g., 0.8% (2/237) [6]; 2.5% (2/79) [31]; 3.23% (2/62) [32]; 4.36% (26/597) [33] and 4.88% (4/82) [34]). These results indicate a higher *E. bieneusi* infection in stray cats in shanghai in this study, posing a public health concern, although it can not be entirely excluded that *E. bieneusi* spores may only pass through the gastrointestinal tract (pseudoparasitism), as identification of *E. bieneusi* DNA from faecal samples is not direct evidence of infection.

**Table 3.** Prevalences of *Enterocytozoon bieneusi* recorded previously in cats and dogs worldwide.

| Host species | Prevalence of *E. bieneusi* % (*E. bieneusi* test-positive sample nos./total tested sample nos.) | Country | Reference |
| --- | --- | --- | --- |
| Cat | 1.40 (2/143) | China | [28] |
| Cat | 2.34 (4/171) | China | [7] |
| Cat | 2.54 (3/118) | Czech Republic | [35] |
| Cat | 3.03 (3/99) | Spain | [42] |
| Cat | 3.33 (2/60) | Brazil | [43] |
| Cat | 3.80 (6/158) | South Korea | [44] |
| Cat | 5.00 (3/60) | Germany | [45] |
| Cat | 5.56 (4/72) | Turkey | [6] |
| Cat | 5.63 (9/160) | China | [9] |
| Cat | 5.77 (3/52) | China | [25] |
| Cat | 6.25 (4/64) | Slovakia | [35] |
| Cat | 6.85 (5/73) | Poland | [35] |
| Cat | 8.33 (1/12) | Switzerland | [46] |
| Cat | 9.09 (4/44) | Poland | [34] |
| Cat | 11.46 (11/96) | China | [47] |
| Cat | 11.63 (20/172) | Australia | [12] |
| Cat | 14.10 (22/156) | China | [48] |
| Cat | 14.29 (1/7) | Japan | [31] |
| Cat | 17.39 (8/46) | Colombia | [49] |
| Cat | 20.31 (79/389) | China | [30] |
| Cat | 31.25 (25/80) | Thailand | [29] |
| Dog | 0.84 (2/237) | Spain | [6] |
| Dog | 2.53 (2/79) | Japan | [31] |
| Dog | 3.23 (2/62) | China | [8] |
| Dog | 4.36 (26/597) | Japan | [33] |
| Dog | 4.88 (4/82) | Poland | [34] |
| Dog | 5.33 (4/75) | Iran | [50] |
| Dog | 5.85 (20/342) | Australia | [12] |
| Dog | 5.98 (29/485) | China | [9] |
| Dog | 6.29 (38/604) | China | [51] |
| Dog | 6.74 (18/267) | China | [25] |
| Dog | 7.69 (2/26) | China | [52] |
| Dog | 8.02 (21/262) | China | [7] |
| Dog | 8.02 (75/935) | Japan | [53] |
| Dog | 8.33 (3/36) | Switzerland | [46] |
| Dog | 8.57 (27/315) | China | [28] |
| Dog | 8.82 (24/272) | China | [10] |
| Dog | 9.59 (7/73) | Spain | [54] |
| Dog | 15.00 (18/120) | Colombia | [55] |
| Dog | 15.52 (54/348) | China | [47] |
| Dog | 18.78 (136/724) | China | [48] |
| Dog | 22.89 (149/651) | China | [30] |

Genotyping studies using previously confirmed *E. bieneusi* samples were excluded.

Here, we took the first step to carry out the association analysis between the risk factor of season and *E. bieneusi* prevalence in cats and dogs. Higher prevalence of *E. bieneusi* was observed in summer, but there was no significant support (*P* > 0.05) (Table 2). Association analysis revealed that stray cats and dogs were significantly associated with higher *E. bieneusi* prevalence than household cats and dogs (*P* < 0.05). Stray cats and dogs had 7.112 times higher risk to infect *E. bieneusi* than that in pet clinics and breeding centers (OR = 7.112; 95% CI [3.263-15.500]) (Table 2). Wang et al. studied *E. bieneusi* from pets and stray cats and dogs in Yunnan in China, and found that stray dogs had higher contaminations of *E. bieneusi* (*P* < 0.05) [7], same as the study of Liu et al. [10]. Similarly, Kváč, et al. conducted *E. bieneusi* investigations from pets and stray cats from three countries (Czech Republic, Poland and Slovakia), and they found that stray cats had higher *E. bieneusi* detection rates than pets [35]. This indicates that stray cats and dogs may have higher risk of *E. bieneusi* infections than that in pets, showing a public health threat. Thus, more studies are needed to monitor this pathogen in stray cats and dogs to prevent the outbreaks of human infections of *E. bieneusi*.

The analysis of ITS sequences data revealed seven *E. bieneusi* genotypes, i.e., CD7, CHN- HD2, D, D-like1, PtEb IX, PtEb IX-like1 and Type IV. Zoonotic genotype D (synonyms: CEbC, Peru9, PigEBITS9, PtEb VI, Peru2, WL8, NCF7, SHW1, MJ10, MJ11, MJ12, isolate 20, ZJR7 and FJS) was the commonest genotype found in humans worldwide, and it was also recorded in 68 animal species in more than 38 countries [23]. Similarly, genotype D was the predominant type in stray cats and dogs in the present study (61.8%; 21/34) (i.e., none of genotype D was found in household cats and dogs in pet clinics and breeding centers in this study), similar to most of other studies (S3 Table). This indicates that stray cats and dogs carrying zoonotic genotype D represent the host reservoirs transmitting *E. bieneusi* from them to humans. Obviously, more studies are needed for further verification.

Genotypes CD7 and PtEb IX found in the present study were commonly found in cats and dogs, also, they were sporadically reported in Bactrian camel, sika deer and white-lipped deer in China [36], whooper swan in China [37] and European badger in Spain [38] (S4 Table). Furthermore, none of these two genotypes had been found in humans yet. The result revealed that these genotypes mainly spread among cats and dogs with occasional dispersal in other animal hosts. Interestingly, genotypes PtEb IX was commonly found in drinking source water, sewer water and wastewater in China [39-41], showing that PtEb IX might be transmissible to susceptible hosts (e.g., cats and dogs) via spore-contaminated water or the environment. However, the exact source and transmission pattern of genotype PtEb IX in cats and dogs are difficult to track. As stated, it is clear that more studies of *E. bieneusi* from humans, other animals and the environment are necessary.

To assess the zoonotic potential of *E. bieneusi* genotypes in the present study, our phylogenetic analysis included ITS sequences of seven genotypes and representatives from ten established *E. bieneusi* Groups (Fig 1). The analysis of these sequence data sets revealed that genotypes CD7, CHN-HD2, PtEb IX and PtEb IX-like1 fall into Group 10, which was mainly reported in cats and dogs, and none of them has been identified in humans yet, showing host specificity. However, genotypes D, D-like1 and Type IV were inferred to be in Group 1 with zoonotic potential (Fig 1). The identification of potentially zoonotic genotypes in cats and dogs in the present study suggested that they might act as host reservoirs transmitting *E. bieneusi* from them to humans, *vice versa*.

### Conclusions

This study recorded *E. bieneusi* in cats and dogs in Shanghai in China. The prevalence of *E. bieneusi* in stray cats and dogs was higher than that in housing cats and dogs, showing that stray cats and dogs have a higher potential to transmit *E. bieneusi* from them to humans, showing a public health threat. The predominant genotype D of *E. bieneusi* identified here in stray cats and dogs have been detected commonly in humans and water samples in other countries, suggesting that stray cats and dogs might act as a reservoir for genotype D that are transmissible to humans. Future studies should elucidate the epidemiology of *E. bieneusi* in humans, animals, water and the environment, in order to provide an informed position on its public health importance in this country. Other studies could be conducted to establish whether some of the genotypes recognised to be potentially zoonotic actually occur in humans in China.

## Acknowledgements

The authors thank colleagues from Shanghai/Minhang Center(s) for Animal Disease Control and Prevention, and staff from pet clinics, breeding centers and shelters for sample collection.

## Authors’ contributions

Sample collection: RSM, YDZ, LMX, HZH, HYG, YH and TM. Designed the study and performed the experiments: YZ and YDZ. Analysis and interpretation: YZ and YDZ. Write the manuscript: YZ. Review the draft: ZGC. All authors read and approved the final version of the manuscript.

## Abbreviations

AIC: Akaike information criteria
BI: Bayesian inference
ITS: Internal transcribed spacer of nuclear ribosomal DNA
*LSU*: Large subunit of nuclear ribosomal DNA gene
MCMC: Monte Carlo Markov Chain
pp: Posterior probability
*SSU*: Small subunit of nuclear ribosomal DNA gene

## Declarations

### Ethics approval and consent to participate

All faecal samples were donated from Shanghai/Minhang Center(s) for Animal Disease Control and Prevention with the consent of their owners or staff. During the whole experimental process, all laboratory work on the study specimens was covered under the Animal Experimental Protocol of Shanghai Veterinary Research Institute (201008): “Use of animal samples for the determination of zoonotic pathogen”.

## Data Availability Statement

All relevant data are within the manuscript and its Supporting Information files.

## Supporting information

**S1 Table.** GenBank accession numbers of all internal transcribed spacer (ITS) of nuclear ribosomal DNA sequences used for phylogenetic analysis (Fig 1), and associated information.

**S2 Table.** Genotypes of *Enterocytozoon bieneusi* characterised from 359 individual faecal samples (sample codes given) from cats and dogs in this study.

**S3 Table.** All *Enterocytozoon bieneusi* genotypes recorded previously in cats (*Felis catus*) and dogs (*Canis familiaris*) worldwide.

**S4 Table.** Genotypes PtEb IX, Type IV, D, CD7 and CHN-HD2 of *Enterocytozoon bieneusi* recorded in different host species and water samples in previous studies. These genotypes were also recorded in the present study.

